# Bacterial and host factors affecting acquisition of *Streptococcus pneumoniae* in a murine model

**DOI:** 10.64898/2026.01.13.699243

**Authors:** Daniel P. Fecko, Cindy Wu, Jeffrey N. Weiser

## Abstract

Initial acquisition is a critical step in the establishment of colonization that in turn enables transmission to new hosts and potentially leads to invasive disease. Here, we studied host and bacterial factors affecting acquisition, as distinct from those affecting colonization density, using *Streptococcus pneumoniae* (Spn) in a mouse model. Acquisition was quantified using nasal inocula of limiting size to determine the proportion of hosts infected, as well as the median infectious dose. Infant mice (<7 days of age) were highly susceptible to acquisition (ID_50_<30 CFU) compared with adults (>6 weeks of age), with rates declining with increasing host age. Prior influenza A infection greatly increased acquisition in adults. Expression of Spn capsule was also an important factor influencing acquisition in adults, with less of a role for the highly susceptible infants. Prior Spn colonization with a heterologous or homologous strain effectively blocked acquisition in infants by completely occupying the upper respiratory tract niche. Immunity due to prior colonization and nasal microbiota were not found to be factors limiting acquisition. Together, our findings show how high carriage rates of encapsulated Spn during early life and in the setting of recent viral infection could be explained by effects on acquisition. Our study describes an approach for studying factors that specifically affect the step of acquisition.

## Introduction

For mucosal pathogens, colonization is the required first step in the pathogenesis of disease (1). *Streptococcus pneumoniae* (Spn) is a prominent member of the nasopharyngeal flora of humans (2, 3). Invasive infections, which are generally a ‘dead end’ for the organism, occur when colonizing bacteria gain access to normally sterile sites, such as the lungs (pneumonia), blood (bacteremia/sepsis), meninges (meningitis), or middle ear space (otitis media) (4). Nasopharyngeal colonization—the carrier state—is particularly common during early life when carriage rates may exceed 50% and generally decline after infancy (2). Accordingly, invasive disease caused by Spn is especially prevalent during early childhood. Rates of disease then decline post-childhood until the later decades of life when a variety of host factors, particularly recent influenza infection, predispose older adults to invasive infection (5, 6). Spn carriage is characterized by sequential or, less commonly, simultaneous events, each lasting from days to months. Isolates of Spn are typically distinguished by antigenic differences among more than 100 unique serotypes (or types) (7, 8). Serotypic variability is due to structural differences in the polysaccharide capsule encasing the bacteria, which is the major virulence determinant of the species and the basis of all licensed pneumococcal vaccines. Polysaccharide vaccines delivered systemically induce high levels of serotype-specific antibody, which is effective at reducing vaccine-type carriage across vaccinated populations (9). It is less clear whether natural carriage generates an adaptive immune response that is sufficient to impact colonization (10, 11). Murine models used by our laboratory recapitulate these key features of natural carriage, including duration of colonization; effect of host age and influenza co-infection; role of capsule; and importance of innate immunity in host defense (12–16).

Colonization is part of a wider process required for mucosal pathogens that is host-to-host transmission (17), which can be broken into three steps. The bacteria first exit via shedding from an original, colonized host. Second, the bacteria are acquired by a new host by entering and attaching to the nasal epithelium. Third, the bacteria replicate and establish colonization in the new host. Using an infant mouse model, we previously investigated bacterial and host factors involved in successful pup-to-pup transmission (18). These studies showed, for example, that influenza co-infection increases secretions that drive shedding in a type I interferon-dependent manner (19). In the absence of viral infection, the mild inflammation induced by the pore-forming Spn toxin, pneumolysin (Ply), is needed for the levels of bacterial shedding permissive for transmission (20). Additionally, we utilized marked strains to characterize population bottlenecks during the Spn life cycle (21). These studies revealed a tight population bottleneck of only a few organisms during the acquisition step, whereas no population bottleneck was detected during shedding or the establishment of colonization. Therefore, it appears that acquisition is the important barrier in the Spn life cycle and, consequently, is also a critical event in Spn pathogenesis. As acquisition has not previously been investigated due to a lack of tractable models, the focus of this report is to take advantage of murine models to define some of the major bacterial and host factors involved.

## Results

### Quantitative assessment of acquisition

To quantify acquisition, intranasal inocula containing serial ten-fold dilutions of Spn (10^1^, 10^2^, 10^3^,… CFU) were compared to estimate the threshold dose required to seed colonization. Because the detection of acquisition depended on a period in which there was bacterial replication and expansion, our primary analysis was based on the proportion of infected hosts (colonized vs. non-colonized), rather than the density of bacteria at an endpoint. We therefore chose 3 days post-inoculation (dpi), when stable colonization is generally achieved, as an arbitrary time point to assess acquisition. This also ensured that the number of cultured Spn exceeded the size of the inoculum and represented colonization rather than retention of the inoculum. A pilot study showed an equivalent level of acquisition at 1 dpi compared to 3 dpi (**Fig. S1**), suggesting that we were not underestimating acquisition because of clearance events earlier than 3 dpi. Colonization was determined by plating of retrograde nasal lavages with a limit of detection of ∼17 CFU/mouse. The proportion of colonized vs. non-colonized mice in each experimental group was then statistically compared using Fisher’s Exact Test. We also estimated the number of organisms required to infect 50% of hosts (ID_50_), which is inversely related to hosts’ susceptibility to acquisition.

### Effect of host age on acquisition

Because infants are highly susceptible hosts, we initiated our study using pups 4-7 days of age. We inoculated with a serotype 23F streptomycin-resistant derivative (23F WT, P2499) of a clinical isolate previously used in experimental human challenge studies where ∼50% of healthy adults became colonized with a dose of 10^3-4^ CFU (Table 1) (22). Spn was acquired in 100% of the pups with each inoculum dose tested (10^1^, 10^2^, 10^3^ CFU) (**Fig. 1A**). In contrast, significantly fewer young adult mice (age 6-8 weeks) became colonized following inoculation with 10^1^ and 10^2^ CFU. Based on these data, we estimated the ID_50_ to be <30 CFU for infants and 129 CFU for young adults. With increasing age in older adult mice, there was a further decrease in acquisition (ID_50_ 318 and 973 CFU at 6 months and 1 year of age, respectively). We also tested a second Spn strain (TIGR4, serotype 4) and found a similar effect of age (infant vs. young adult) on acquisition (**Fig. 1B**). Thus, by using infant mice we could assess factors that decrease acquisition, whereas in adult mice, we could examine factors that increase acquisition.

**Figure 1.**
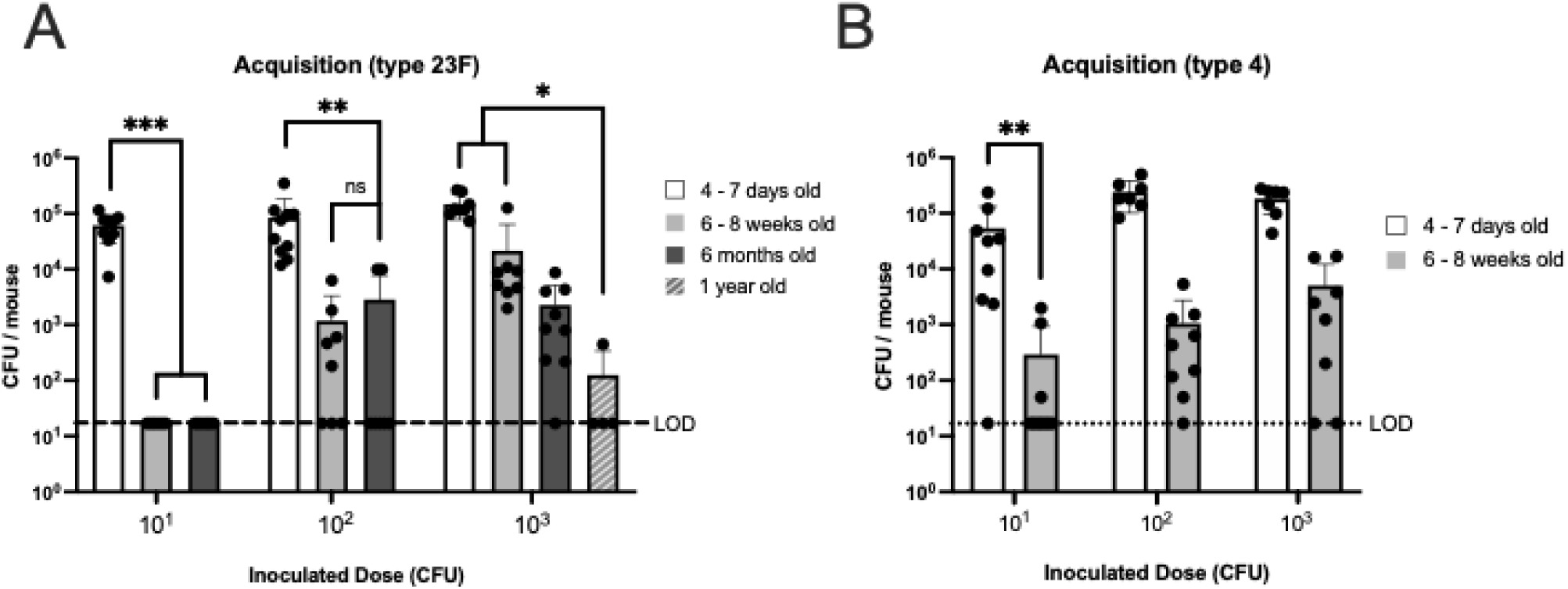
Effect of age on susceptibility to pneumococcal acquisition. **A.** Acquisition of *Streptococcus pneumoniae* (Spn) type 23F WT **B.** Acquisition of Spn TIGR4. Mice of the indicated age were inoculated with a dose of 10^1^, 10^2^, or 10^3^ CFU per mouse and colony-forming units (CFU) determined in retrotracheal lavages at 3 days post-inoculation. Statistical significance determined using Fisher’s exact test comparing number of mice with at least 17 CFU/mouse to those that did not show any colonies above that limit of detection (LOD). Each data point represents an individual animal. Horizontal bars denote the mean ± standard deviation of the group. We considered *p* < 0.05 to be statistically significant or included the *p* value instead of asterisks. **, *p* < 0.01, ***, *p* < 0.001.

**Table 1:** Bacterial strains.

| Strain | Strain Number | Description | Reference |
| --- | --- | --- | --- |
| 23F WT | P2499 | Streptomycin-resistant derivative of P1121(type 23F) | Shen et al. (2019) (40) |
| TIGR4 | P2942 | Novobiocin-resistant derivative of TIGR4 (type 4) | Ma et al. (2025) (47) |
| 23F $\Delta cps$ | P2109 | P1121 with Janus cassette deleting the *cps* locus, kanamycin resistant | Hamaguchi et al. (2018) (48) |
| 23F (4) | P2140 | P2109 with TIGR4 (type 4) *cps* locus inserted, spectinomycin resistant | Lysenko et al. (2010) (49) |
| 23F $\Delta ply$ | P1726 | P2499 with clean deletion of *ply*, streptomycin resistant | Shen et al. (2019) (40) |
| 23F $\Delta cbpD, cibAB$ | P2576 | P2499 with clean deletion of *cbpD* and Janus cassette replacing *cibABC*, kanamycin resistant | Shen et al (2019) (40) |
| 23F $\Delta blp$ | P2700 | P2499 with Janus cassette replacing *blp* locus, kanamycin resistant | Aggarwal et al (2023) (50) |

To test whether *in vivo* adaptation enhanced acquisition, we directly inoculated nasal lavages obtained from colonized mice into adults. Direct inoculation, however, was no more effective than *in vitro*-grown organisms of the same dose (**Fig. S2**).

### Influenza A infection increases pneumococcal acquisition

Because Spn colonization and disease are more common clinically in the setting of recent flu infection, we determined whether influenza A virus (IAV) affects acquisition using our adult mouse model. The mouse-adapted strain H3N2 IAV x31 or a PBS-vehicle control was administered intranasally prior to inoculation with 23F WT. IAV co-infection significantly increased the acquisition rate when delivered either 3 or 7 days before Spn inoculation, with the ID_50_ in adults reduced to <15 and <18 CFU, respectively, compared to the PBS control ID_50_ of 1389 CFU (**Fig. 2A**). The timing between the two agents coincides with high susceptibility in human patients to post- influenza Spn pneumonia. IAV titers were measured in lavages and were only positive in the group given x31 3 days before bacterial challenge (**Fig. 2B**). Thus, the impact of recent flu infection on Spn acquisition was maintained for at least 7 days following the bacterial challenge, even though viral co-infection had cleared by the time of lavage 10 days post-IAV-infection.

**Figure 2.**
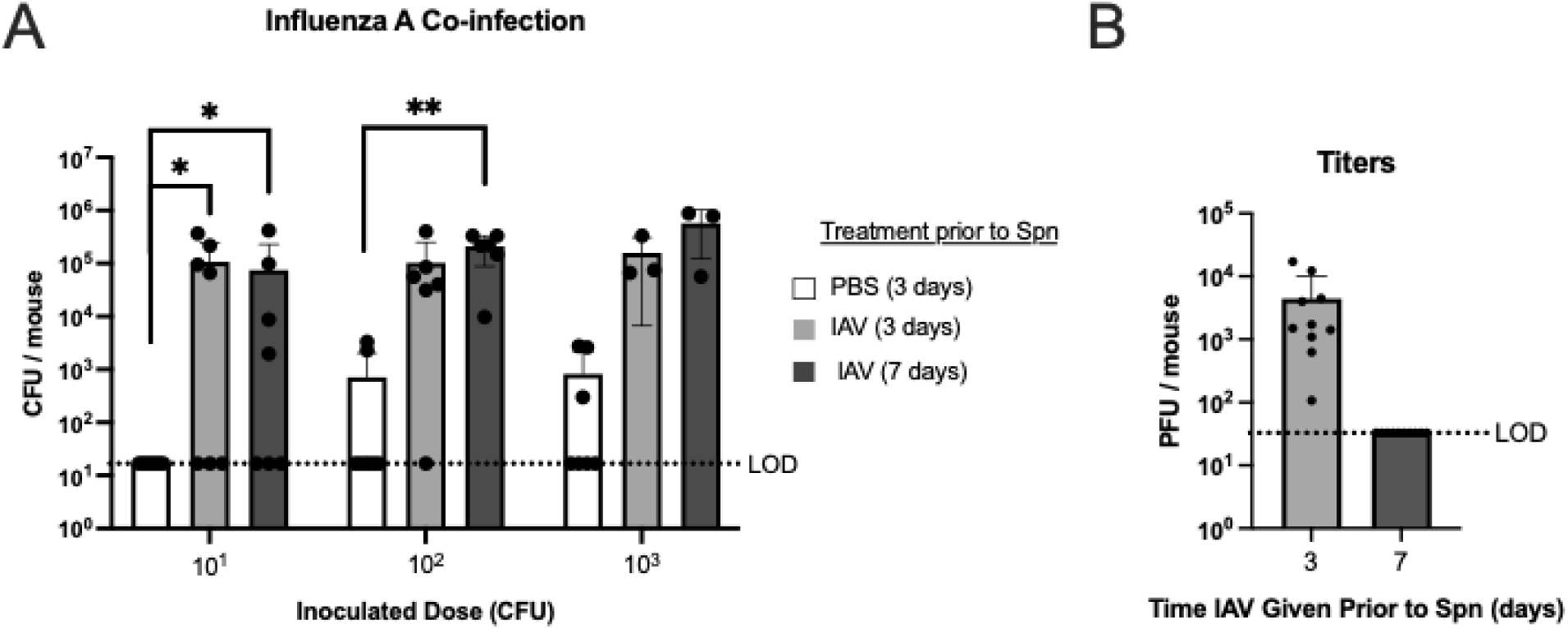
Effect of influenza A co-infection on susceptibility to pneumococcal acquisition. **A-B**. Adult mice were inoculated IN with 20,000 PFU of Influenza A virus (IAV) strain x31 either 3 or 7 days before Spn inoculation. PBS-control was given 3 days before Spn inoculation. Colony-forming units (CFU) were determined in retrotracheal lavages at 3 days post-Spn inoculation with a dose of 10^1^, 10^2^, or 10^3^ type 23F WT Spn/mouse. **A**. Acquisition of Spn in IAV co-infected mice. Statistical significance determined using Fisher’s exact test comparing number of mice with at least 17 CFU/mouse to those that did not show any colonies above the limit of detection (LOD). **B.** Plaque-forming units (PFU) of x31 Influenza A virus in RT lavages of mice in panel A. Each data point in the figures represents an individual animal. Horizontal bars denote the mean ± standard deviation of the group. LOD, dashed line. *, *p* < 0.05, **, *p* < 0.01.

### Effect of the nasal microbiota on acquisition

The nasal mucosa contains microbiota that increase in diversity with host age (23). During colonization, Spn likely competes with existing microbiota for nutrients and physical space in the host. To test the contribution of the flora to limiting acquisition, adult mice were treated for one week with high-dose streptomycin added to their drinking water prior to 23F WT challenge. This broadly active antibiotic was selected because acquisition was determined using a streptomycin-resistant Spn strain (23F WT). As expected, this treatment reduced the number of non-Spn microbes measured in nasal lavages by standard culture methods (**Fig. 3A**). However, there was no detectable difference in the rate of acquisition in mice treated with streptomycin drinking water, suggesting a minimal role for the microbiota in acquisition success (**Fig. 3B**).

**Figure 3.**
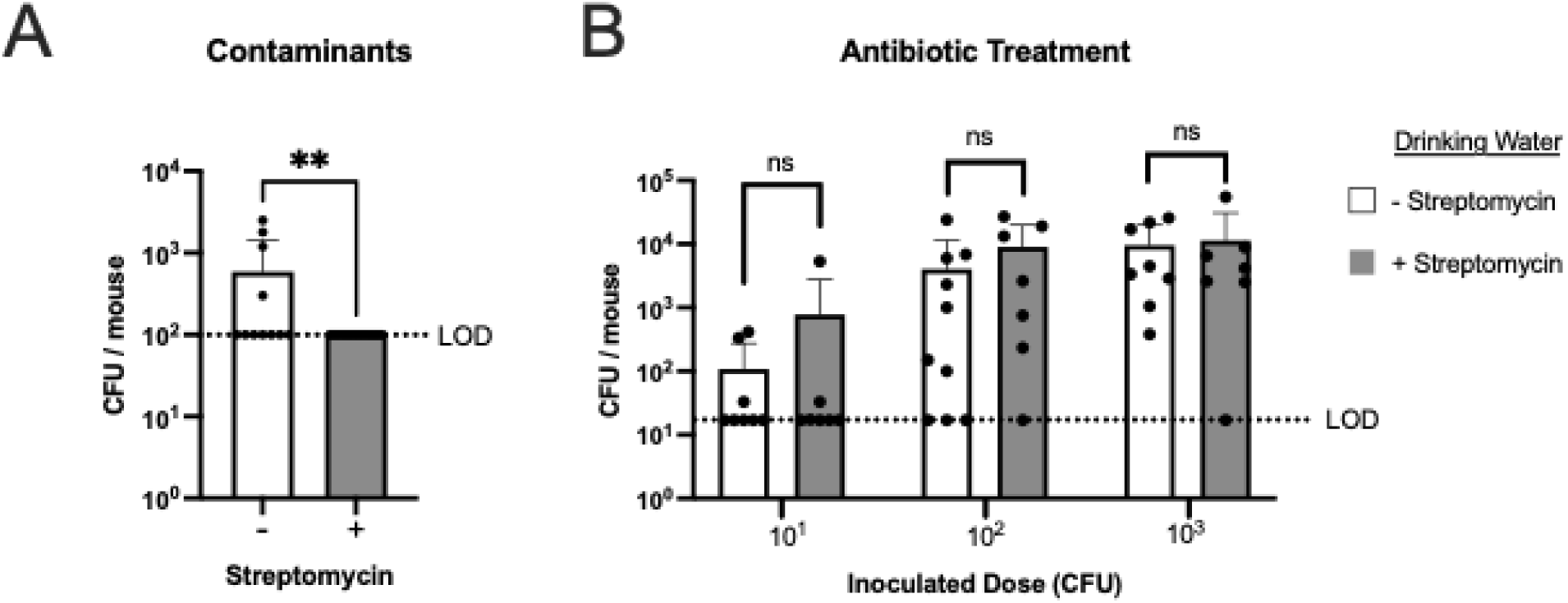
Antibiotic depletion of the microbiome treatment does not influence pneumococcal acquisition. **A-B.** Adult mice were given streptomycin in the drinking water starting (500 mg/mL) a week before Spn inoculation and continuing until euthanasia. **A.** CFU of non-Spn in RT lavages 3 days after Spn inoculation. Statistical significance determined using Mann-Whitney test. Limit of detection (LOD) of contaminants is 100 CFU/mouse. **B.** Acquisition of type 23F WT (streptomycin-resistant) Spn. Colony-forming units (CFU) were determined in retrotracheal lavages at 3 days post-Spn inoculation with a dose of 10^1^, 10^2^, or 10^3^ Spn/mouse. Statistical significance determined using Fisher’s exact test comparing number of mice with at least 17 CFU/mouse to those that did not show any colonies above the LOD (dashed line). Each data point in the figures represents an individual animal. Horizontal bars denote the mean ± standard deviation of the group. **, *p* < 0.01, ns, not significant.

### Bacterial factors in acquisition

Because expression of a capsule increases Spn colonization density, we examined its contribution to acquisition(15). An isogenic, unencapsulated mutant of 23F WT, 23FΔ*cps*, was not acquired by adult mice at doses of 10^1^-10^3^ CFU (**Fig. 4A**). The ID_50_ of this construct was 5.9×10^4^ CFU. Restoration of capsule expression (construct expressing a type 4 capsule) corrected the deficit (ID_50_ = 3.8×10^2^ CFU), confirming the importance of capsule in acquisition in adult mice. In contrast, capsule did not affect Spn acquisition in the more highly susceptible infant mice (**Fig. 4B**). (ID_50_ of the unencapsulated mutant is <15 CFU, similar to that of the parent and capsule-restored strains.) A caveat of the 23FΔ*cps* colonization after large doses is that the low density detected could represent retention of the inoculum rather than acquisition leading to replication and colonization.

**Figure 4.**
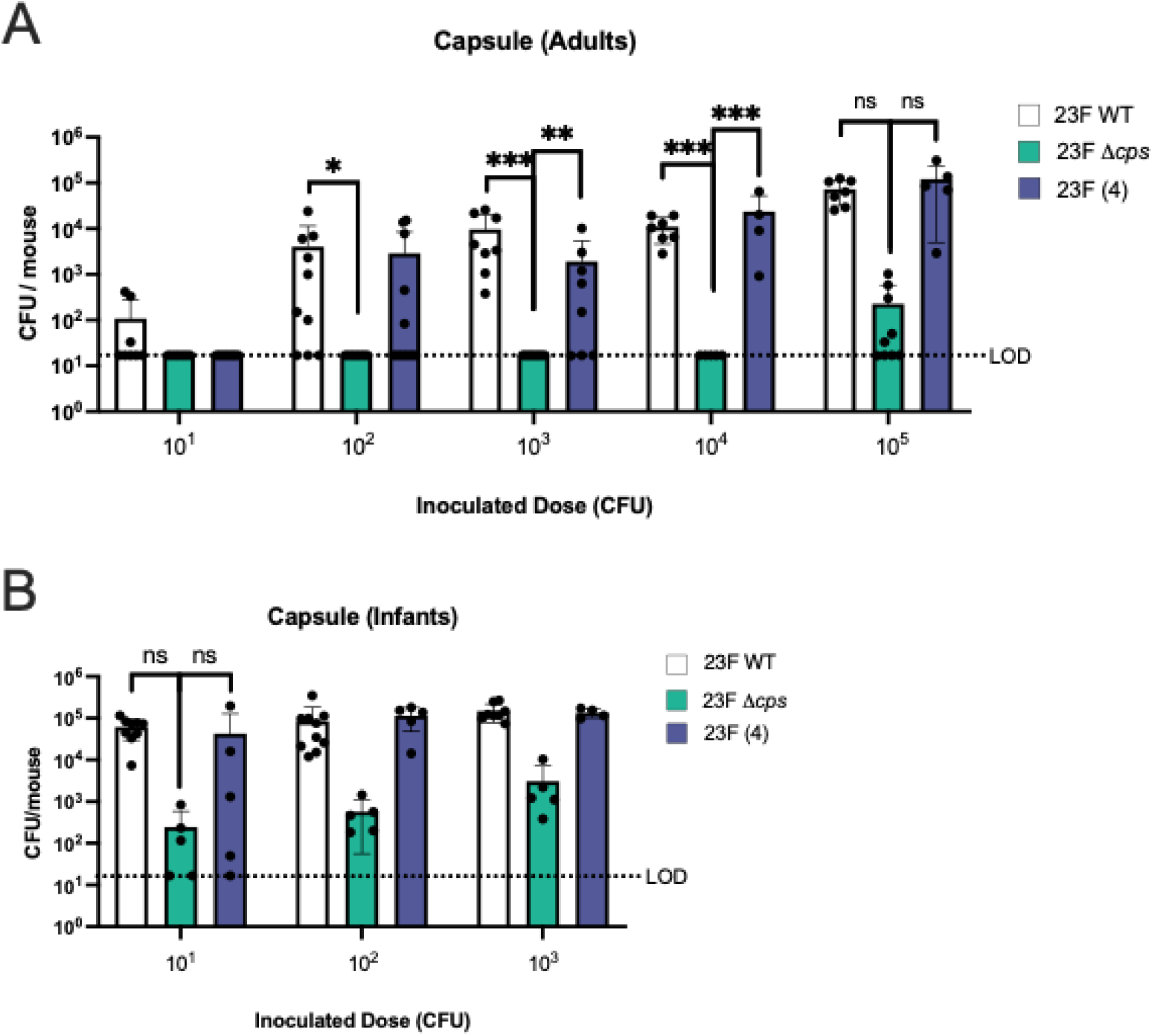
Effect of capsule on pneumococcal acquisition. **A.** Acquisition in adult mice (6-8 weeks old). **B.** Acquisition in infant mice (4-7 days old). CFU of Spn 23F WT, capsule knockout (Δ*cps*), or capsule-restored construct expressing the type 4 capsule (23F (4)) in RT lavages 3 days post-Spn-inoculation with the dose indicated below the graphs. Statistical significance determined using Fisher’s exact test comparing number of mice with at least 17 CFU/mouse to those that did not show any colonies above the limit of detection (dashed line). Each data point in the figures represents an individual animal. Horizontal bars denote the mean ± standard deviation. *, *p* < 0.05, **, *p* < 0.01, ***, *p* < 0.001, ns, not significant.

Another major determinant of Spn-host interaction is the pore-forming toxin pneumolysin, which could alter the early inflammatory milieu and thereby affect acquisition. Some non-hemolytic Spn, such as certain lineages of serotype 1, are uncommonly carried, perhaps due to low acquisition (24). However, there was no effect on acquisition in adults from deletion of the *ply* gene (**Fig. S3**).

### Pre-existing pneumococcal colonization impacts acquisition

Next, we tested whether pre-established colonization inhibited acquisition. Infant mice were first colonized with 23F WT (or given PBS without Spn) and challenged 24 hours later with varying doses of strain TIGR4 to quantify its acquisition. As expected, 23F WT consistently colonized at a high density (>10^5^ CFU/ml). Pre-colonization with 23F WT was associated with a significant reduction in the rate of acquisition of TIGR4 and a corresponding increase in its ID_50_ (1.5×10^3^ CFU vs. <15 CFU in the PBS control group) (**Fig. 5A**). This effect was dependent on pre-colonization, since simultaneous challenge with an equal mixture of the two strains resulted in efficient acquisition of both (**Fig. 5B**). This was the case despite the fact that the 23F WT strain outcompeted the TIGR4 at higher inocula (**Fig. 5C**). Moreover, even when pre-colonized infants were challenged with a homologous strain (distinguished by different resistance markers), acquisition was significantly attenuated (**Fig. 5D**).

**Figure 5.**
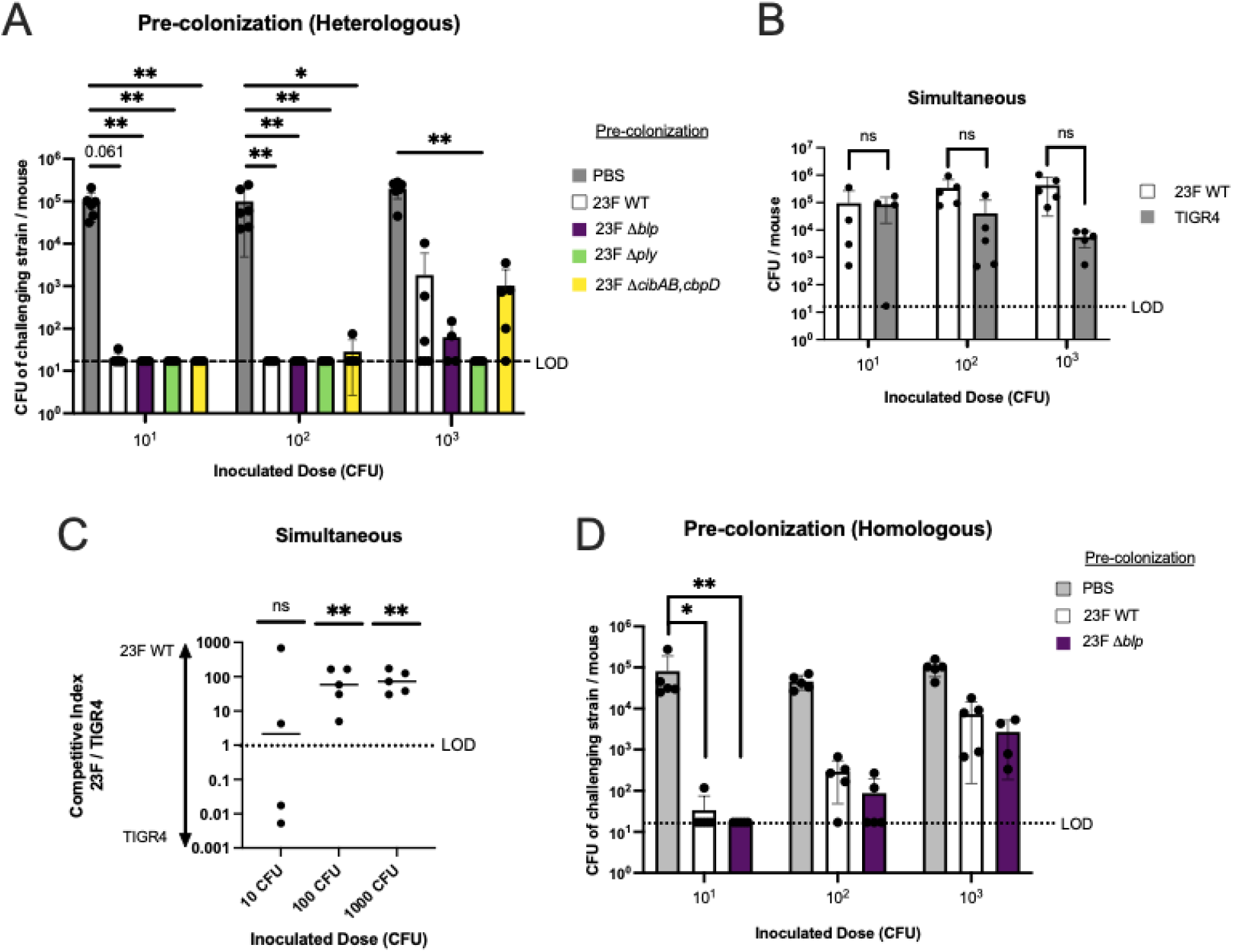
Pre-existing pneumococcal colonization inhibits acquisition. **A.** Acquisition in infants colonized sequentially with heterologous strains. Infant mice were pre-colonized Spn type 23F strains: wildtype (WT), *blp* locus knockout (*Δblp*), pneumolysin knockout (Δ*ply*), fratricide knockout (Δ*cibAB*,*cbpD*), or a PBS control. One day later these mice were inoculated with Spn TIGR4 (challenge strain) at 10^1^, 10^2^, or 10^3^ Spn/mouse. CFU of Spn TIGR4 was determined in RT lavages after 3 days. **B.** Acquisition in infants colonized simultaneously with heterologous strains. Infant mice were inoculated with the same dose of 23F WT and TIGR4 at 10^1^, 10^2^, or 10^3^Spn/mouse together and then the CFU for each strain determined after 3 days. **C.** Competitive index of 23F WT to TIGR4 for each mouse tested in panel B. **D.** Acquisition in infants colonized sequentially with homologous strains. Infants were first pre-colonized with Spn type 23F strains: wildtype (WT), *blp* locus knockout (*Δblp*), or a PBS control, then one day later inoculated with isogenic Spn 23F WT (challenge strain) containing a different antibiotic-resistance marker. Mice were inoculated with the challenge strain at a dose of 10^1^, 10^2^, or 10^3^ CFU/mouse and CFU of the challenge strain determined in RT lavages 3 days later. Statistical significance determined using Fisher’s exact test comparing number of mice with at least 17 CFU/mouse to those that did not show any colonies above the limit of detection (dashed line). Each data point in the figures represents an individual animal. Horizontal bars denote the mean ± standard deviation. *, *p* < 0.05, **, p < 0.01, ns, not significant.

To explore the mechanism involved in pre-established colonization impeding acquisition, we tested several possibilities. These included a competitive advantage mediated by *blp*-locus-expressing bacteriocins or the expression of fratricins (CibAB and CbpD). We also considered the effect of pneumolysin-mediated inflammation induced by the pre-colonizing strain. Each mutant colonized at similarly high levels (**Fig. S4, S5)**. Pre-colonization using isogenic constructs (*Δblp*; Δ*cibAB* and Δ*cbpD*; and Δ*ply*) of 23F WT had no difference from pre-colonization by 23F WT in the rate of acquisition of TIGR4 (**Fig. 5A**). Rather, our data suggests a ‘winner takes all’ model whereby the size of the Spn colonization niche is limited and once fully occupied acquisition of an incoming challenger is greatly restricted. This inhibitory effect of the pre-colonizing strain can only be partially overcome when the challenger is given at a very high dose (Total CFU of pre-colonized strain plus challenger strain in the lavage remains the same across all doses, i.e. fully occupied niche (**Fig. S6**).

### Limited role of immunity from prior pneumococcal colonization on acquisition

As natural infection is characterized by multiple carriage events with different strains, we examined whether immunity from prior colonization affects acquisition. Mice were intranasally colonized with 23F WT at 21 days of age and left undisturbed until 9 weeks of age to ensure that colonization with the ‘immunizing strain’ was essentially cleared, serving as an “immunizing” process. These mucosally immunized adults were then challenged with 10^4^ CFU of a strain of the same serotype and genetic background. At this dose, there was no decrease in the rate of acquisition compared to the ‘PBS immunization’ control group (**Fig. 6A**). A larger inoculum compared to previously described experiments was used, because of the variability in assay controls when lower doses were tested. Thus, we could not exclude the possibility of differences in acquisition using lower inocula. A 23F WT strain whole-cell serum IgG ELISA was used to confirm an immune response in mice colonized with the immunizing strain (**Fig. 6B**).

**Figure 6.**
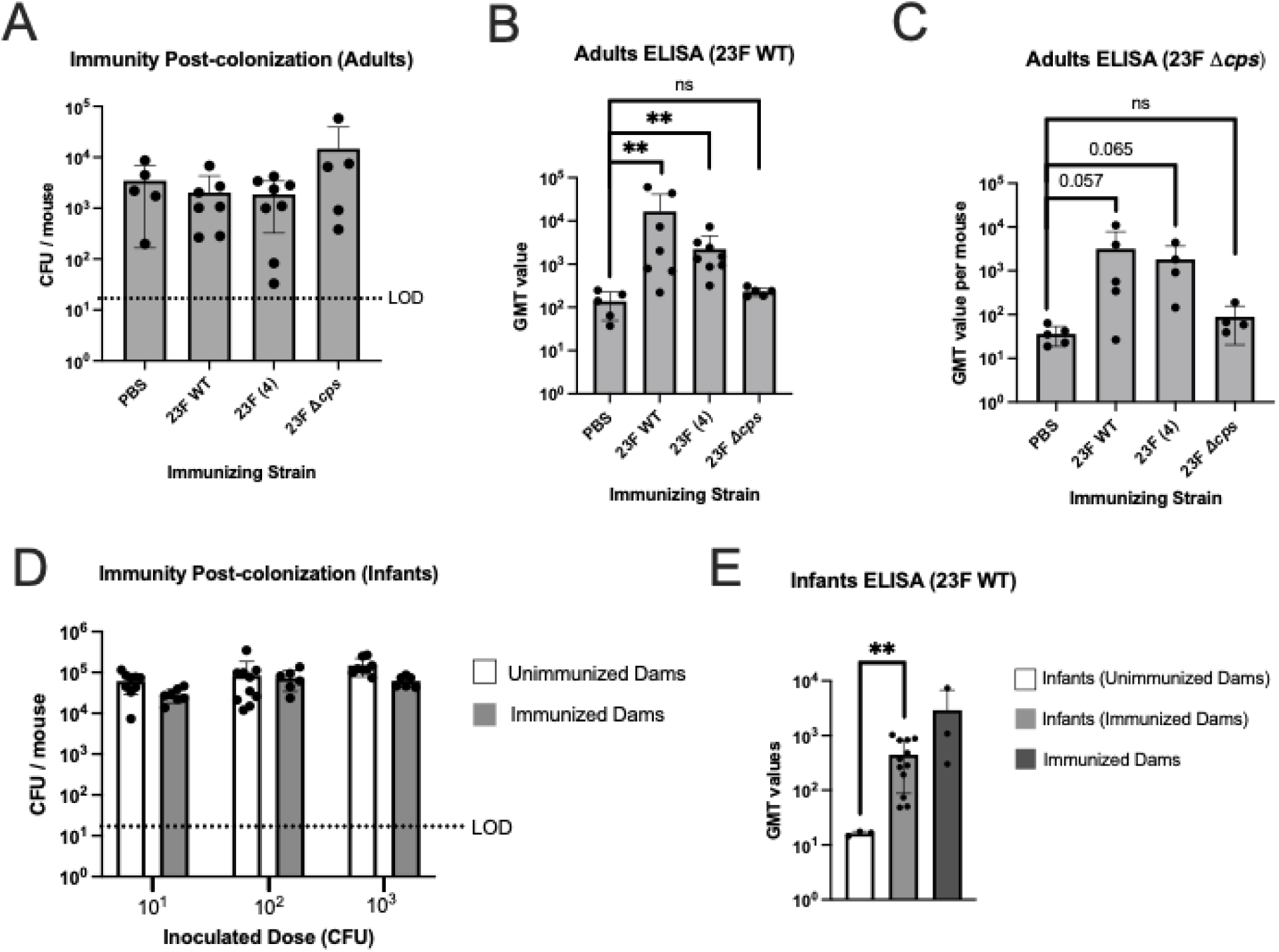
Immunity from prior pneumococcal colonization is not sufficient to block acquisition. **A-C.** Mice were ‘immunized’ at 21 days of age with PBS control or colonization with Spn strains: 23F WT, 23F genetic background expressing the type 4 capsule 23F (4), or 23F genetic background with capsule locus deleted (23F Δ*cps*). Six weeks later following clearance mice were inoculated with 10^4^ CFU of Spn 23F WT and CFU of Spn type 23F WT determined in RT lavages 3 days later. **A**. Effect of immunity from prior colonization on acquisition. **B.** GMT values for serum IgG in a whole-cell ELISA with Spn type 23F WT for mice shown in panel A. **C.** GMT values for serum IgG in a whole-cell ELISA with Spn type 23F capsule locus deleted (23F Δ*cps*) for mice shown in panel A. **D-E.** Female mice were ‘immunized’ at 21 days of age by colonization with Spn type 23F, then 6-8 weeks later gave birth to pups. These pups were inoculated at 4-7 days old with 10^1^, 10^2^, or 10^3^ CFU of Spn 23F WT and housed continually with the dams. Infants were euthanized and RT lavaged 3 days after inoculation. **D.** Acquisition in infants born to immunized or unimmunized dams. **E.** GMT values for serum IgG in a whole-cell ELISA with Spn type 23F WT for the infants in panel D and their dams. Statistical significance determined using Fisher’s exact test comparing number of mice with at least 17 CFU/mouse to those that did not show any colonies above the limit of detection (dashed line). Each data point in the figures represents an individual animal. Horizontal bars denote the mean ± standard deviation. We considered p < 0.05 to be statistically significant or included the p value instead of asterisks. **, p < 0.01, ns, not significant.

We also immunized mice using a construct of 23F WT expressing the type 4 capsule (capsule-switch strain) and then challenged them with the 23F WT strain to isolate an effect of serotype-independent immunity. Detectable levels of serum antibody were measured using the 23F WT whole cell ELISA, suggesting that cross-reactive immunity was generated to non-capsular antigens (**Fig. 6B**). This was also confirmed using a whole-cell ELISA to detect reactivity against an isogenic, unencapsulated mutant (**Fig. 6C**). Immunization with the capsule-switch mutant also failed to block acquisition. Finally, we tested an isogenic, unencapsulated mutant as the immunizing strain, but this failed to induce significant levels of cross-reactive antibody or decrease acquisition of the 23F WT parent—a result that could be due to low levels of colonization by the immunizing strain (**Figs. 4A, 6A, 6B and 6C**).

While we could not similarly test the direct effect of mucosal immunity during infancy because of the time required for clearance, we examined vertically transferred immunity from dam to pup. We tested acquisition in pups 4-7 days of age born to dams immunized by prior colonization with the same strain. There was, however, no difference in the rate of acquisition between pups born to previously colonized and those of non-colonized dams (**Fig. 6D**). We confirmed that pups from immunized dams had cross-reactive antibody at the time of sacrifice using the 23F WT whole-cell ELISA (**Fig. 6E**).

Together these findings suggest that acquired immunity from prior colonization, including that against both capsular and non-capsular antigens, does not have a major impact on the rate of acquisition.

## Discussion

Our investigation focused on the main bacterial and host factors linked to carriage prevalence to better understand the contribution of acquisition to overall colonization dynamics. The power of our approach comes from using limiting dilutions to estimate the 50% infectious dose, which differs from the general practice of infecting with large inocula.

The study of Spn acquisition shown in this murine model has also been examined in the experimental human challenge of adult volunteers. Using the same parent strain, experimental human colonization of healthy adults showed an ID_50_ of ∼10^3^ CFU, only one log higher than we found in our far smaller murine hosts (22). The human challenge study, therefore, provides an important validation of our approach that relied on a murine model. The animal model, although less physiologically relevant, is more tractable and allows for the testing of a range of factors that cannot be readily examined in the human model.

Successful acquisition requires the retention of organisms that first encounter the mucosal surface. Within the first 30 minutes following inoculation, Spn is found embedded in the firm mucus layer overlying the epithelial surfaces of the nasopharynx (14). Spn expresses a number of exoglycosidases that alter mucus characteristics to promote tighter mucus interactions and enhance colonization (25). An additional requirement of successful acquisition is the evasion of mechanical clearance by mucociliary flow of the loose mucus layer that sweeps the upper respiratory tract (URT). Without the repulsive effects of its capsule, unencapsulated Spn is quickly entrapped in the luminal mucus and cleared (14). This could account for the acquisition defect of unencapsulated Spn observed in our adult mice model and the lower prevalence of natural nonencapsulated carriage isolates in humans (26). The interplay of Spn with mucus is also affected by host age. Infants have less efficient mucociliary clearance in the URT (27). Enhanced retention due to this deficit could explain why we found a lesser requirement of encapsulation for acquisition in infant mice.

A number of bacterial surface adhesins that promote Spn binding to epithelial cells have been described (4). As acquisition is the first step leading to carriage, these would be expected to also contribute to colonization. A previously described Tn-seq library screen, however, failed to demonstrate a non-redundant effect of known adhesins on Spn’s ability to colonize infant mice, and so these were not individually examined in our study (28).

One of the major findings of this study was the age-related effect on acquisition with a ∼50-fold increase in susceptibility (based on the ID_50_) of infant mice compared to young adults, with a gradual decrease with age thereafter. Differences in the mucus layer in infants facilitates closer proximity of Spn to the epithelial surface (27). Infants also have lower levels of the major antimicrobials of airway surface fluid, including lysozyme and lactoferrin, which could be factors that make the infant URT more permissive to initial events in colonization (29). Importantly, increased acquisition by infants correlates with increased transmission among pups when tested using a mouse model (30).

Similarly, increased acquisition following recent IAV infection could be due to altered mucus flow and mucus retention caused by impaired mucociliary clearance (31). Although IAV interferes with early innate immunity to Spn and increases capillary leakage and nutrient availability, these factors are more likely to affect the step of replication rather than acquisition (32, 33). In a study of experimental human Spn carriage, evidence of recent asymptomatic respiratory syncytial virus or rhinoviruses was associated with Spn acquisition (34). This study, which was timed to avoid flu season, suggests that the effects of recent viral infection on acquisition could extend to other respiratory viruses. The effect of influenza A on acquisition described here correlates with increased transmission reported in both ferret and mouse models (35, 36).

We considered the possibility that the URT microbiome could have a negative impact on acquisition because of interspecies interactions or inflammatory effects on the host environment. Depletion of URT flora prior to challenge, however, had no effect on acquisition in adult mice. On the other hand, an experimental human Spn carriage study in healthy adults did find a positive association between acquisition and a more diverse microbiome prior to inoculation (37). In a separate study, immediate nasal clearance was associated with increased numbers of mucosal neutrophils, suggesting that a more inflammatory milieu, which could be affected by the microbiota as well as host age, is a determining factor in acquisition (38, 39). Our inbred mice housed under highly controlled pathogen-free environmental conditions are unlikely to be impacted by high levels of endogenous inflammation. In this regard, we tested whether pneumolysin-mediated inflammation impacts the ability of a strain to be acquired but observed no effect.

Our data showed that acquisition could be blocked by pre-existing, dense colonization with another Spn strain, suggesting a competitive advantage for the resident over a newcomer. Our observation for acquisition resembles the ‘owner-intruder’ population dynamic that leads to a ‘winner takes all’ scenario previously proposed for Spn colonization (40). Our investigation, which used heterologous strains, did not indicate a role for inflammation triggered by the resident (tested with a *ply* mutant) or Blp-bacteriocin-mediated Spn-Spn competition in affecting acquisition. We also observed no contribution of anti-Spn fratricins, controlled by the competence (Com) system for niche dominance, affecting acquisition of unrelated strains. We had previously described fratricin-mediated niche dominance for isogenic strains during colonization (41). Instead, our results were consistent with a limited overall URT niche size for Spn that, once full, excludes newcomers in a dose-dependent manner. It is unclear what restricts this ceiling for colonization density, although certain factors such as IAV co-infection raise its limit (compare density of adult colonization in Figs. 1A and 2A). It is possible that the availability of nutrients, which are released in greater quantity during influenza-mediated inflammation, is the cause of the limited niche size. Our data also raises the question of how natural co-colonization with more than a single strain occurs. We propose that this becomes possible once the density of the resident drops below the threshold that excludes newcomers.

It is not surprising that immunity from prior colonization fails to inhibit subsequent acquisition events, considering that sequential carriage of multiple strains is common clinically (10). Our observations show that even prior colonization with an isogenic strain, which generates humoral immunity to both capsular and non-capsular antigens, is not sufficient to inhibit acquisition. A caveat of our analysis is that an immune response was confirmed by demonstrating the presence of serum antibody, because low antibody levels and their dilution in the lavage make mucosal antibody difficult to measure.

Our results contrasted with human experimental challenge data, in which 0 out of 10 volunteers were recolonized with the same strain 3 months to a year after first challenge, compared to a rate of 60% for volunteers challenged for the first time with a similar dose. In this human study, however, the first challenge was most likely boosting existing immunity from previous natural pneumococcal exposure, whereas our naïve mice had no exposure prior to immunization. Passive URT immunization with human monoclonal type-specific IgA1 was previously shown to block acquisition through its agglutinating activity but only if not cleaved by the IgA1-protease expressed by Spn (42). Agglutination by anti-capsular IgG, which is IgA1-protease insensitive, is also associated with protection against experimental human Spn carriage (43). This human challenge model has also demonstrated that serotype-specific memory B cells, but not levels of circulating IgG, at the time of exposure are associated with protection against acquisition (44). In fact, serotype-specific serum IgG, which accesses mucosal surfaces if present in sufficient quantity as occurs following the administration of PCV vaccine, has been shown to inhibit acquisition of experimental human carriage in a double-blind randomized control trial (45). The level of IgG generated by PCV immunization, however, generally far exceeds that induced by a colonization event tested here in mice. Another consideration is that blocking acquisition of only a few organisms may require far fewer specific antibodies than clearing an already-established colonization. Prior carriage, however, does not appear to generate sufficient antibody to impact acquisition. In summary, our analyses of bacterial and host factors demonstrate the importance of host age, influenza infection, capsule expression, and prior colonization status in affecting Spn acquisition. The effect of these factors on the key step of acquisition could explain why natural carriage of encapsulated Spn is highest during infancy and increased in the setting of recent viral infection, as well as why simultaneous carriage events are less common than expected by rates of single carriage (46).

## Methods

### Spn strains and growth conditions

Spn strains used in this study are listed in Table 1. Spn was grown statically in tryptic soy broth (TSB) at 37°C in a water bath until reaching an OD of 1 at 620 nm, measured by the Spectronic 200 (Thermo Scientific). Spn were then spun down at 10,000 g, washed once with sterile PBS, spun down again and resuspended with sterile PBS, then diluted to desired CFU/ml. To count CFU, Spn were plated in tenfold dilutions in PBS onto tryptic soy agar (TSA) plates containing catalase (6,300 U/plate) and incubated overnight in 5% CO_2_ at 37°C.

### Mice

All animal experiments followed the guidelines summarized by the National Science Foundation Animal Welfare Act (AWA) and the Public Health Service Policy on the Humane Care and Use of Laboratory Animals. The Institutional Animal Care and Use Committee (IACUC) at New York University Grossman School of Medicine oversees the welfare, well-being, proper care and use of all animals, and they have approved the protocol used in this study (IA16-00538).

Wild-type C57BL/6J (strain 00664) were obtained from Jackson Laboratory. All mice were bred and handled in our research facility. Water and a standard rodent diet were provided ad libitum. Male and female mice were both used. Pups were housed with a dam (mother) for the entire duration of the study. *S. pneumoniae* colonization did not impact the weight gain of mice. Where indicated streptomycin (500 mg/L) was added to the drinking water for one week prior to Spn challenge and continuing until euthanasia 3 days after inoculation. Contaminants in nasal lavages were estimated by quantitative culture using tryptic soy 5% sheep blood agar plates incubated under microaerophilic conditions at 37°C for 16 hours in 5% CO_2_. Spn distinguished by colony phenotype and confirmed by optochin sensitivity were excluded from the quantification of contaminants.

### Acquisition Model

Spn was inoculated intranasally into mice in sterile PBS with a pipette tip, without anesthesia. Infants (4-7 days old) were inoculated using 3 μL and adults (6-8 weeks) with 20 μL. One or three days later we euthanized the mice via CO_2_ asphyxiation followed by cardiac puncture. We performed a retrotracheal (RT) lavage with sterile PBS (500 ml) using a 30-gauge needle for infants and a 26-gauge needle for adults, inserting the needle into the trachea and collecting the liquid exiting from the nares. The lavage was then plated in ten-fold dilutions on an TSA-catalase plate supplemented by antibiotics specific to the colonizing strain(s) (250 μg/ml kanamycin, 200 μg/ml streptomycin, 200 μg/ml spectinomycin, 5 μg/ml novobiocin) and colonies enumerated after growing in an incubator overnight in 37°C for 16 hours or 64 hours at 30°C in 5% CO_2_.

For IAV co-infection, 20,000 PFU of IAV/HKx31 was administered intranasally into mice, and then either 3 days or 7 days later, Spn was inoculated. Mice were euthanized and lavaged 3 days after Spn 23F WT inoculation. Viral titers were determined in MDCK cells by standard plaque assay in the presence of TPCK (tosylsulfonyl phenylalanyl chloromethyl ketone)-treated trypsin (Thermo Scientific) (19).

Where specified, mice were inoculated with 10^3^ CFU of a pre-colonizing strain 1 day before the challenging strain, with euthanasia and lavage 3 days after the challenging strain. Both the pre-colonizing strain and challenge strain were plated for quantitative culture as described above.

For direct inoculation, 10^3^ CFU of Spn was inoculated into infants 4-7 days old, then 3 days later euthanized and lavaged as described above. The lavage fluid was then used to inoculated adult mice (6-8 weeks old) either undiluted or at the 10^-1^ dilution, corresponding to 10^3^ and 10^2^ CFU, respectively. Infant lavages were also plated to confirm the density of Spn.

Under all conditions described above, transmission between co-housed animals was <10%.

### Colonization-Induced Immunity

To test the effect of immunity following previous colonization, mice were inoculated at 21 days of age with 4 x 10^5^ CFU of an immunizing strain. Sentinel mice were euthanized and lavages cultured to test that the immunizing strain had cleared from the URT or was present at a very low density (<10^2^ CFU/ml). The remainder of the mice were then challenged with 10^4^ CFU 23F WT Spn 6 weeks later (9 weeks old) and euthanized and lavaged 3 days after 23F WT inoculation, with blood collected via cardiac puncture. Lavages were plated on both streptomycin-supplemented plates to quantify acquisition of 23F WT, as well as on an antibiotic-containing plates specific for the immunizing strain to confirm loss of or minimal residual colonization.

To test the effect of passive immunity transferred from dam to pup, female mice were inoculated with 23F WT at 21 days old and 3 weeks later paired for breeding with naive male mice. Pups were inoculated with 23F WT at 4-7 days old and euthanized and lavaged 3 days later, with blood collected by cardiac puncture.

### Enzyme-Linked Immunosorbent Assay

Spn-specific IgG following colonization was measured by an ELISA. Spn strains were grown to 1.0 OD at 620 nm, diluted 1:10 in sterile PBS, and 100 μL of the dilution was allowed to incubate overnight in 4°C in 96-well Immulon 2HB plates (ThermoFisher 3455). Wells were washed 3 times with Brij 35 solution (0.05% Brij 35 in PBS) and then blocked with 1% BSA/PBS (Fisher 9048-46-8) for 1 hour at RT. Blood was obtained via cardiac puncture, allowed to clot, and spun down using SST tubes (BD Microtainer) to acquire serum. Serum from mouse cardiac punctures were added in serial dilutions in 1% BSA/PBS and allowed to incubate overnight at 4°C. After washing with the Brij 35 solution, secondary anti-IgG alkaline-phosphatase-conjugated antibodies (Sigma A3688 057K6016) were added at 1:4000 dilution in 1% BSA/PBS and incubated for 1.5 hours. Wells were washed, then 1 mg/mL pNPP (sera-care 5120-0057) in diethanolamine (Thermo 34064) was added and incubated for 1 hour. Fluorescence was detected at 415 nm using the SpectraMax M3 (Molecular Devices).

### Statistical Methods

Acquisition of Spn was compared for each inoculum dose between two experimental groups using Fisher’s exact test. Where appropriate, we estimated the weighted average dose required to achieve a rate of 50% infection (ID_50_) using the Reed-Muench method.

Comparisons of ELISA antibody titers between two groups used the Mann-Whitney U test or Student’s t-tests for normally distributed data. A Mann-Whitney test comparing the ratio of competing strains to an identical number of hypothetical values of 1 was used to assess the significance of competitive indices. GraphPad Prism version 10.6.1 was used for statistical analyses.

## Acknowledgements

The authors thank Dr. Stacie Barlett for assistance with viral titers.

## Author Contributions

D.P.F. led the project, designed and performed experiments, analyzed data and wrote the manuscript. C.W. began the project, designed and performed experiments, and analyzed data. J.N.W. directed the project, designed experiments, analyzed data and wrote the manuscript.

## Supplemental Figures

**Figure S1.**
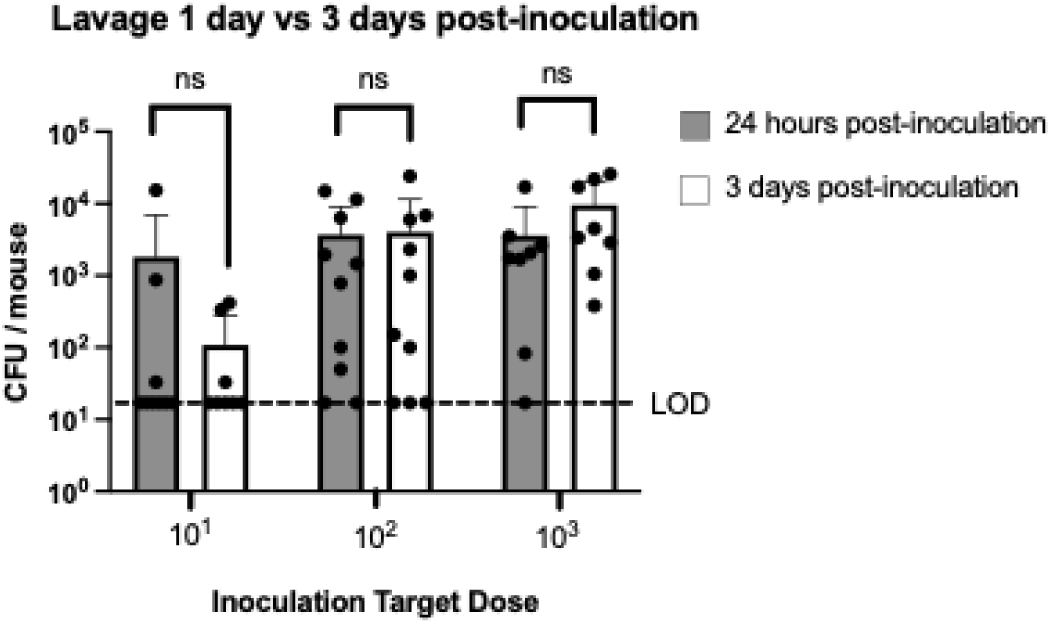
Effect of amount of time after challenge on rate of acquisition. Acquisition of Spn 23F WT in lavage when mice 6-8 weeks old were euthanized and lavaged either 1 day or 3 days post-inoculation. Mice were inoculated with a dose of 10^1^, 10^2^, or 10^3^ CFU/mouse. Statistical significance determined using Fisher’s exact test comparing number of mice with at least 17 CFU/mouse to those that did not show any colonies above that limit of detection (dashed line). Each data point in the figures represents an individual animal. Horizontal bars denote the mean ± standard deviation. ns, not significant.

**Figure S2.**
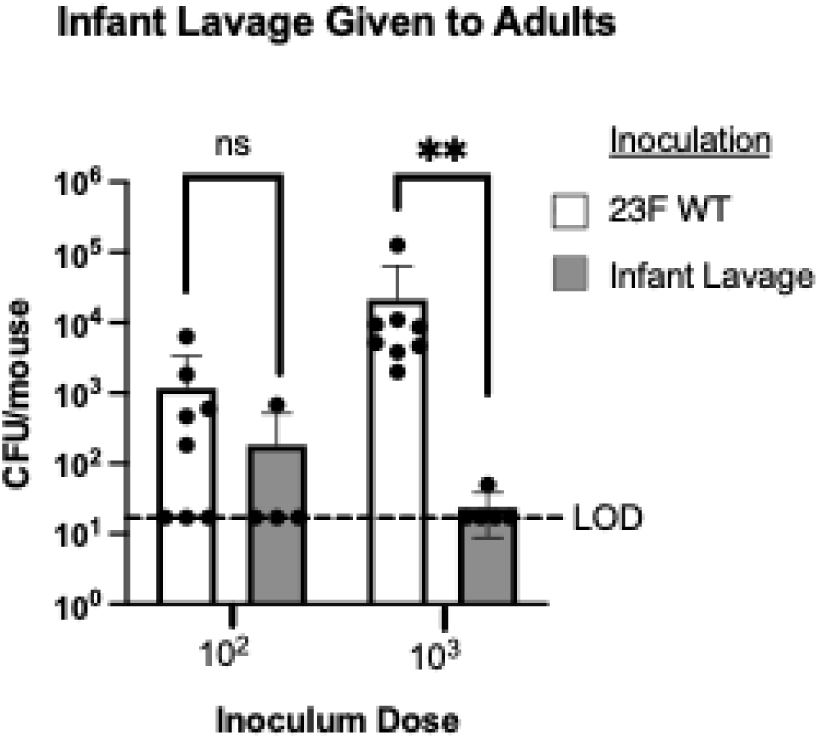
In vivo propagation does not improve acquisition. Infant mice 4-7 days old were inoculated with Spn 23F WT and then lavaged 3 days later. This lavage was directly inoculated into adult mice 6-8 weeks old either undiluted or at the 10^-1^ dilution, corresponding to inocula of 10^3^ and 10^2^ CFU/mouse, respectively. Infant lavages were plated to check correct CFU. The inoculated adults were then euthanized and lavaged 3 days post-inoculation. Results were compared to inoculation with in vitro grown 23F WT. Statistical significance determined using Fisher’s exact test comparing number of mice with at least 17 CFU/mouse to those that did not show any colonies above that limit of detection (dashed line). Each data point in the figures represents an individual animal. Horizontal bars denote the mean ± standard deviation. **, *p* <0.01. ns, not significant.

**Figure S3.**
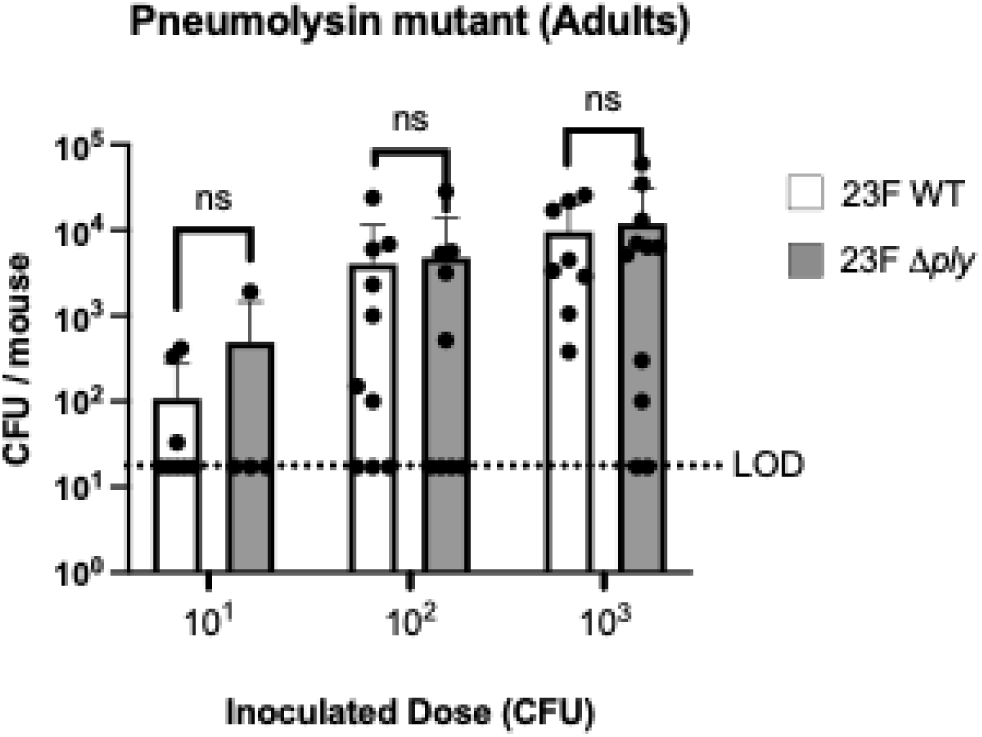
Knocking out pneumolysin does not affect acquisition. Acquisition of 23F WT compared to 23F strain with a pneumolysin deletion mutant (Δ*ply*) in adult mice 6-8 weeks old, lavaged 3 days after inoculation. Mice were inoculated with a dose of 10^1^, 10^2^, or 10^3^ CFU/mouse. Statistical significance determined using Fisher’s exact test comparing number of mice with at least 17 CFU/mouse to those that did not show any colonies above that limit of detection (dashed line). Each data point in the figures represents an individual animal. Horizontal bars denote the mean ± standard deviation. ns, not significant.

**Figure S4.**
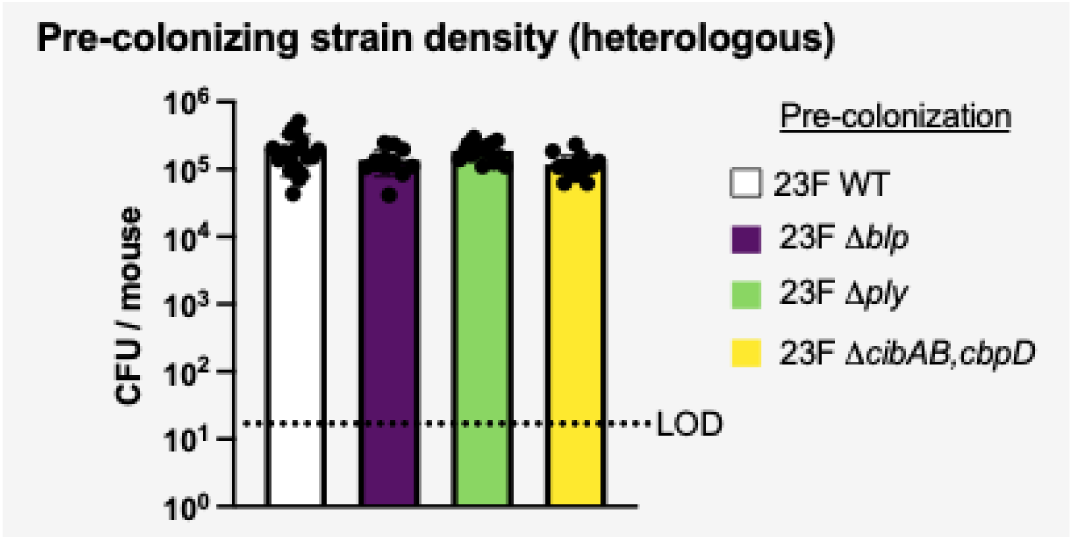
Equivalent colonization level for pre-colonizing strains in heterologous challenge. CFU of Spn in lavages of infant mice 3 days after challenge strain inoculation. Mice in Fig 5A. were pre-colonized with 10^3^ CFU of various Spn 23F strains (wildtype (WT), *blp* locus knockout (*Δblp*), pneumolysin knockout (Δ*ply*), fratricide knockout (Δ*cibAB*,*cbpD*), and then one day later challenged with 10^1^, 10^2^, or 10^3^ CFU doses of TIGR4. CFU of pre-colonizing strain in RT lavages shown.

**Figure S5.**
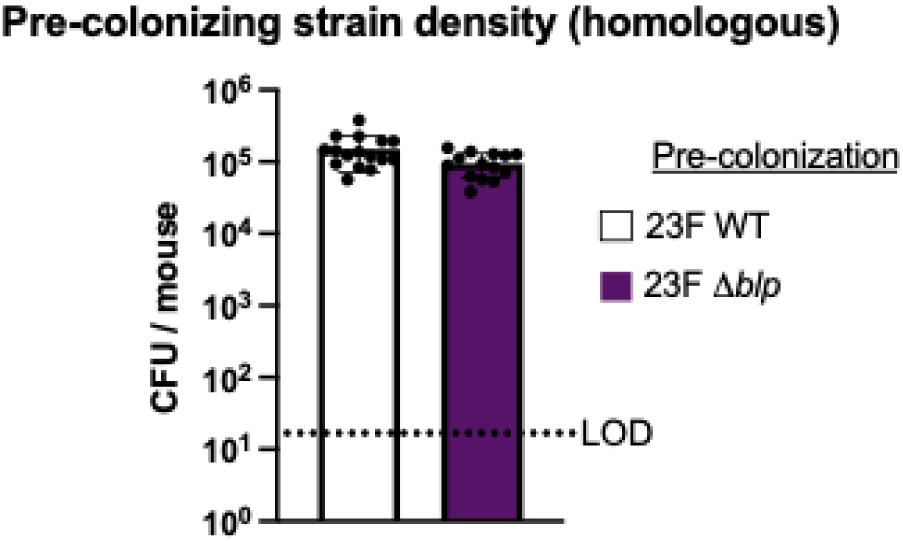
Equivalent colonization level for pre-colonizing strains in homologous challenge. CFU of Spn in lavages of infant mice 3 days after challenge strain inoculation. Mice in Fig 5D. were pre-colonized with 10^3^ CFU of a Spn 23F strain (23F WT or 23F *blp* mutant (*Δblp*)) and one day later challenged with Spn 23F WT. 3 days post-challenge these mice were euthanized, lavaged, and the lavage was plated with 2 different antibiotics to distinguish strains. CFU of pre-colonizing strain in RT lavages shown.

**Figure S6.**
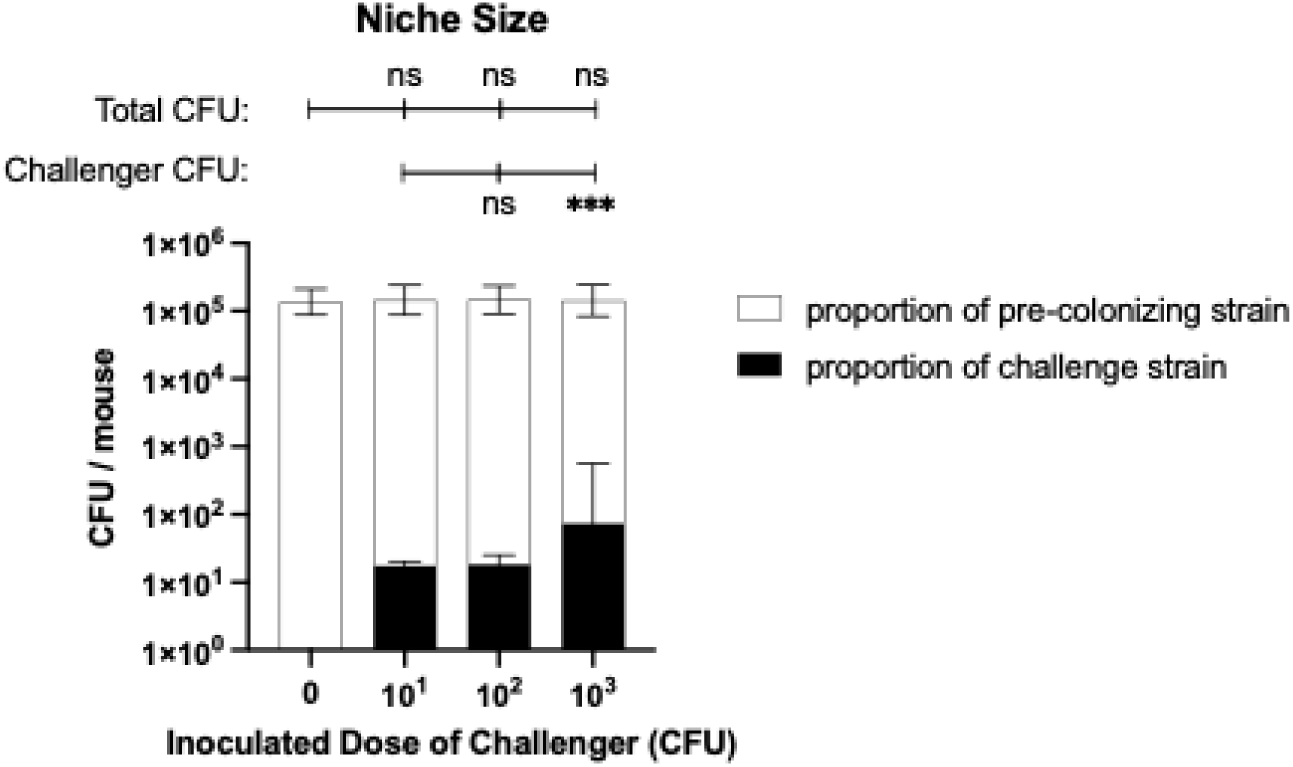
Competition for a limited colonization niche size. Mice in Fig, 5A were pre-colonized with various Spn 23F strains and then challenged one day later with 10^1^, 10^2^ or 10^3^ CFU of TIGR4. The different pre-colonizing strains are grouped together as “pre-colonizing strain” and all mice are combined based on the size of the challenge strain dose. Above: Total CFU (pre-colonizing plus challenge strain) was compared to group without pre-colonization (0). CFU of the pre-colonizing strain was also compared to the group receiving the 10^1^ inoculum of the challenge strain. Horizontal bars denote the mean ± standard deviation. ***, *p* <0.001. ns, not significant.

